# Degradability tunes ECM stress relaxation and cellular mechanics

**DOI:** 10.1101/2024.07.28.605514

**Authors:** Badri Narayanan Narasimhan, Stephanie I. Fraley

## Abstract

In native extracellular matrices (ECM), cells can use matrix metalloproteinases (MMPs) to degrade and remodel their surroundings. Likewise, synthetic matrices have been engineered to facilitate MMP-mediated cleavage that enables cell spreading, migration, and interactions. However, the intersection of matrix degradability and mechanical properties has not been fully considered. We hypothesized that immediate mechanical changes result from the action of MMPs on the ECM and that these changes are sensed by cells. Using atomic force microscopy (AFM) to measure cell-scale mechanical properties, we find that both fibrillar collagen and synthetic degradable matrices exhibit enhanced stress relaxation after MMP exposure. Cells respond to these relaxation differences by altering their spreading and focal adhesions. We demonstrate that stress relaxation can be tuned through the rational design of matrix degradability. These findings establish a fundamental link between matrix degradability and stress relaxation, which may impact a range of biological applications.

**Table of contents:** This work reveals that matrix degradability, through its effects on stress relaxation, is an important cellular mechanotransduction cue. Cell-scale mechanical characterization shows that collagen gels and degradable synthetic gels display enhanced stress relaxation post-degradation. Stress relaxation is then tuned by systematically varying degradability, resulting in the regulation of cell spreading. This identifies degradability as a key chemomechanical design feature.

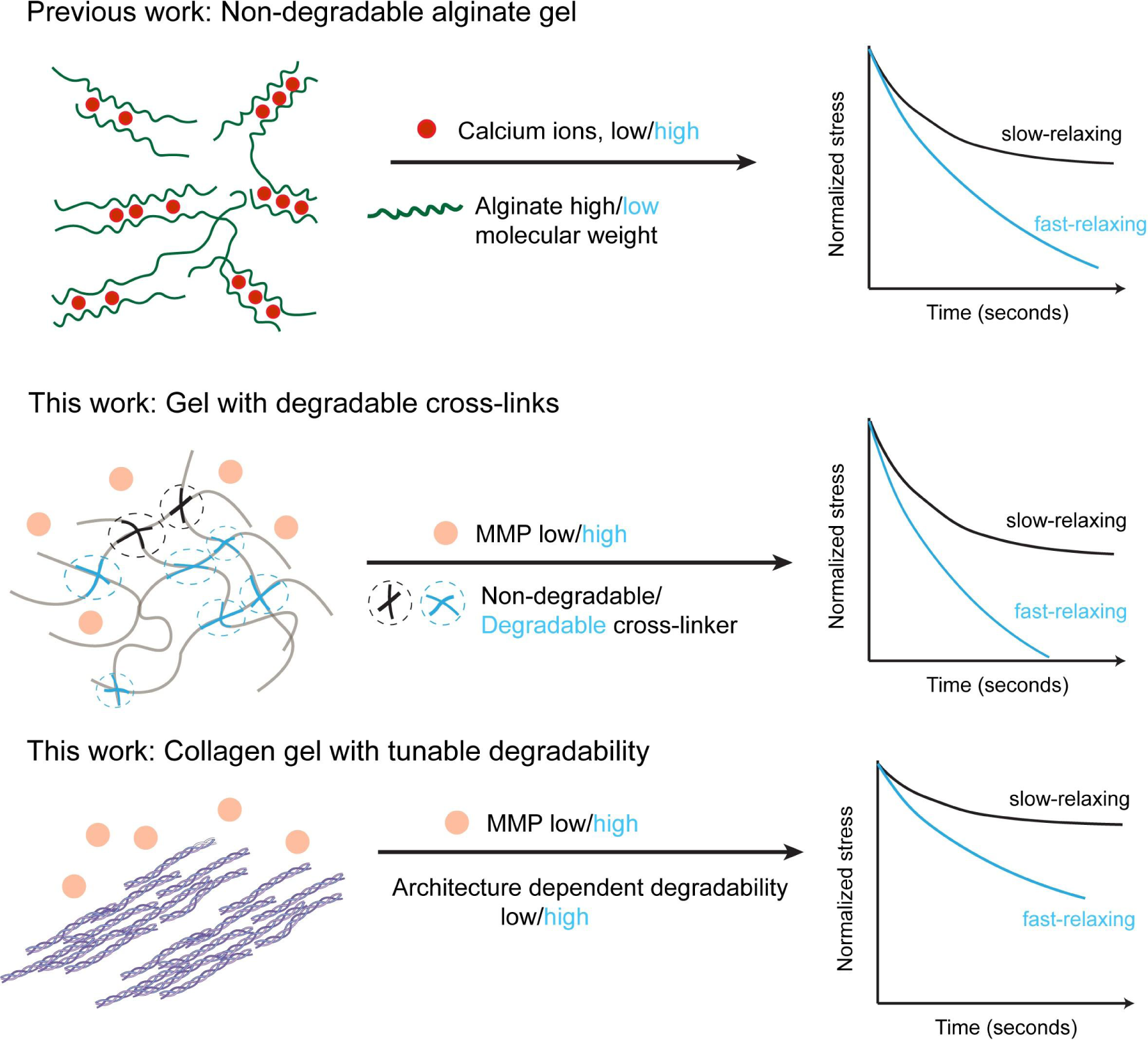

## 1 Introduction

Native ECM is a complex material composed of a variety of proteins that defines key mechanical and biochemical properties of tissues. The development of tunable synthetic biomaterials has advanced our understanding of how cells sense and respond to specific mechanical features of the ECM. A variety of crosslinking mechanisms have been used to systematically vary the elastic modulus, which has revealed the influence of stiffness on cell differentiation, adhesion, proliferation, and gene expression^[1–3]^. Independently, viscoelasticity has been tuned through distinct crosslinking strategies or with interpenetrating networks of polymers, and this has also revealed impacts on cell behavior and communication, influencing cell adhesion, alignment, migration, and remodeling of the local ECM^[4–6]^. These studies defined the importance of static mechanical environments in controlling cellular processes. However, the mechanical properties of native ECM can be dynamic, with different timescales and types of changes corresponding to different biological processes.

*In vivo*, much of the ECM dynamics are controlled by cellular remodeling. Cells remodel their environment either by degrading it with MMPs or by depositing and reorganizing matrix proteins^[7,8]^. The degradability of the ECM has been shown to significantly influence cell behavior, and several studies suggest that degradation of ECM is a prerequisite for successful adhesion and force-transmission from the cell to the ECM^[9,10]^. ECM degradability has also been shown to affect stem cell differentiation fate, cell protein deposition, and even the transition of cells from single to collective modes of migration^[11–13]^. Here, degradability is defined as the rate at which the cells digest the matrix. In 3D systems, degradability also relates to the time evolution of space created by cellular MMPs that allows cells to change shape, grow, and move. Thus, cells tend to spread better in degradable gels than the non-degradable gels due to the physical confinement imposed by non-degradable surroundings^[14]^. Perhaps because of its more obvious role in 3D confinement, not much work has looked at the role that degradation rate may play on 2D cell behavior. However, in one study, Peng et al^[15]^ found that human mesenchymal stem cells seeded on soft gels with higher degradability surprisingly promoted osteogenesis over adipogenesis, a process that normally requires a stiff or stress-relaxing viscoelastic matrix^[16]^. This could indicate that cells experience local changes in tension as a function of time on matrices with higher degradability, but this has not yet been explicitly proven. However, the current paradigm is that degradability and mechanical properties are uncoupled^[17]^. Few studies on elastic degradable gels and assessment of remodeling by fibroblasts revealed that, while there are changes in viscoelasticity of the environment^[18,19]^, the contribution from degradation versus matrix deposition could not be separated. It remains unclear what cells sense in the degradable environment after degradation by MMPs and how it regulates cellular tension.

In native ECM, collagen is the most abundant protein present and much work has been dedicated to understanding how it is degraded and remodeled due to the importance of this process in development, regeneration, and several diseases^[20]^. Unlike most synthetic ECM systems, collagen is composed of three polypeptide chains that assemble into a triple helical structure^[21]^, and these assemble into higher order structures called fibrils that are typically sub micron in diameter and present a distinct 67 nm periodicity^[22]^. Many such collagen fibrils then associate to form fiber structures present in tissues. At the fibril scale, MMPs specifically cleave at locations that are ¾ of the length from the N-terminus.^[23]^. This action can result in unwinding of the hierarchical structure and further cleavage^[24]^. Depending on the extent of degradation, this can result in breaks in the cross-links of collagen and microstructural alterations either through changes in collagen organization or by reducing the thickness of collagen fibrils ^[25]^. Yet, an understanding of how these changes impact fibrillar network mechanics is still lacking.

The breaking of crosslinks, whether in synthetic polymer gels or collagen fibril networks, can increase polymer/fibril entanglements. Plastic flow could subsequently result which can enhance the stress relaxation properties. Therefore, we hypothesized that cells may mechanically sense local ECM degradation through its effects on stress relaxation, a dynamic time-dependent mechanical property. To test this, we produced native and synthetic hydrogels of varying degradability, treated them with collagenases, and assessed their post degradation mechanics and cellular responses. First, we found that fibroblasts responded differently to the collagenase treated collagen matrices as a function of their degradability. The fibroblast response to collagen matrices suggested that stress relaxation could be the determinant of the cellular response post-degradation. Then, we set out to measure the changes in the mechanical properties of the matrices post degradation. For this, we used colloidal probe atomic force microscopy (AFM) based force relaxation measurements to assess the relevant cell-scale mechanical properties. Indeed, stress relaxation differences correlated with cellular response. To isolate degradability as a tunable feature, we then designed synthetic hydrogels with MMP-degradable cross-linkers. To decouple the impact of matrix stiffness, we rationally altered the ratio of MMP-degradable to non-degradable cross-links in the gels so that their post-degradation stiffness was similar to their non-degradable counterparts. AFM relaxation experiments revealed changes in stress relaxation after degradation on the degradable gels but not on the non-degradable gels. Fibroblasts cultured on the synthetic gels after degradation by collagenases were assessed for cell spreading and focal adhesions. Notably, we were able to replicate the relationship between degradability vs. stress relaxation similar to collagen gels. On gels of equivalent stiffness but different degradability (and therefore stress relaxation), fibroblasts spread to different extents and had different numbers of focal adhesions. Furthermore, we found that these responses are cell-type dependent. These findings highlight the importance of degradability-mediated stress relaxation, which can be an important mechanical design parameter in future biomaterials.

## 2 Results

### 2.1 Collagen degradability tunes cell spreading in response to collagenase

Our main goal was to understand whether and how differences in degradability translate into differences in cell mechanosensing. To begin, we used our previously demonstrated macromolecular crowding method to produce collagen gel substrates with different degradability while maintaining the same density ^[12]^ (Figure 1A). We used 2 mg/mL and 8 mg/mL polyethylene glycol (PEG) as the crowding agent with 2.5 mg/mL collagen to produce more degradable (MD) and less degradable (LD) gels, respectively. The PEG was removed by washing after gelation ^[12]^. The degradation rate of MD and LD gels was assessed by measuring the absorbance at 313 nm after collagenase incubation, which confirmed that MD gels exhibited a higher degradation rate than the LD gels (Figure 1B). Then, we treated the gels with a low concentration of collagenase (10 µg/mL) to mimic enzymatic degradation by cells. The corresponding collagenase-treated gels were denoted as MDx and LDx gels. These more degradable and less degradable collagen substrates with and without collagenase treatment serve as our initial model system.

**Figure 1.**
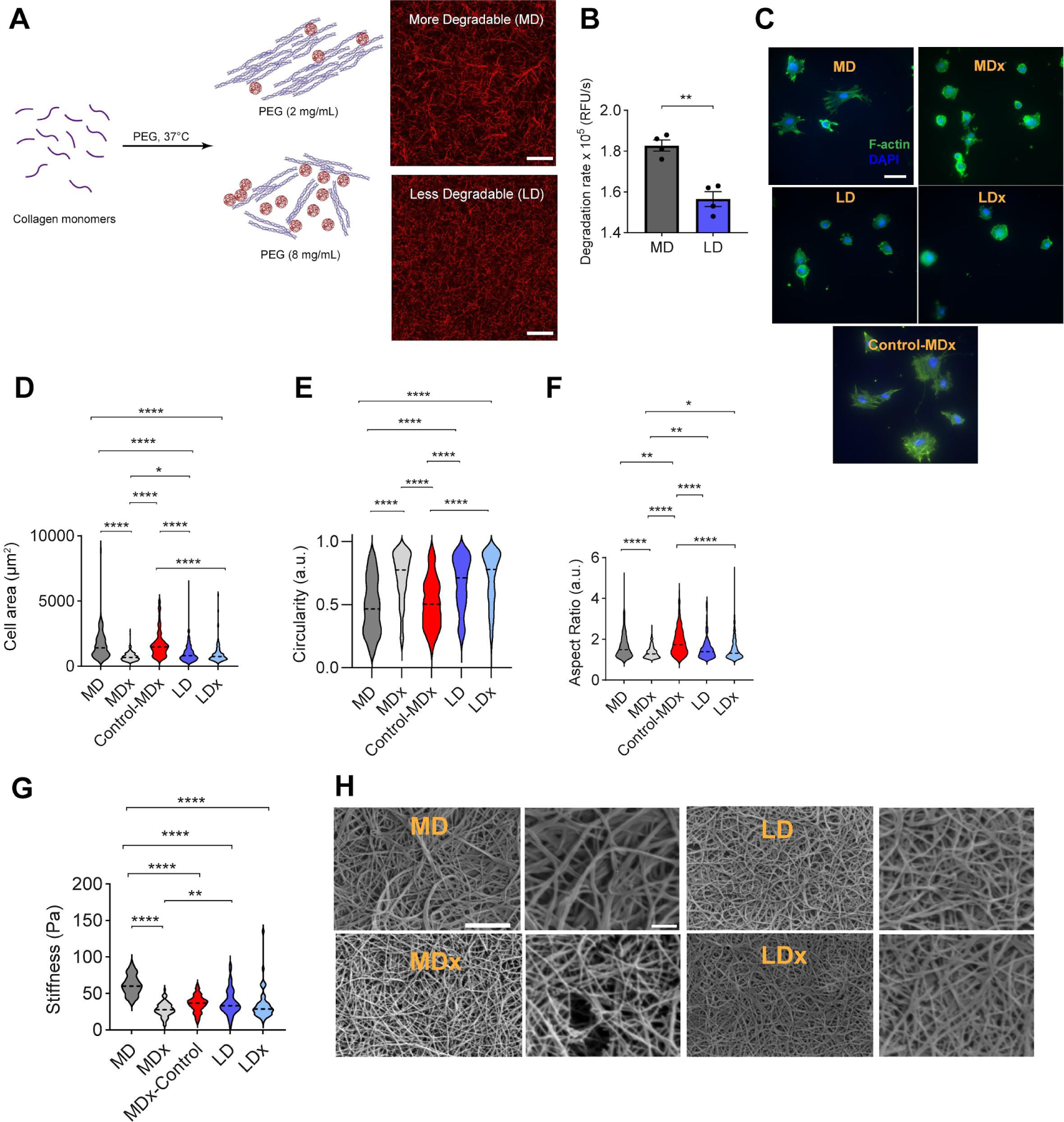
Cells sense changes in collagen architecture and degradability independent of stiffness. MD - More degradable, MDx - More degradable gels with collagenase treatment, LD - Less degradable, LDx - Less degradable gels with collagenase treatment, MDx-Control - same stiffness as that of MDx gels. (A) Scheme for producing collagen gels of varying degradability through macromolecular crowding with PEG (created with BioRender.com). Scale bar represents 20 µm. (B) Degradation rate quantified by monitoring change in absorbance at 313 nm after adding collagenase, Student’s t-test was performed between samples. (C) Representative immunofluorescent images of HFFs spreading on MD, MDx, LD and LDx gels. F-actin in green, nucleus in blue. Scale bar represents 50 µm. Quantification of HFF spreading: (D) Area, (E) Circularity, and (F) Aspect ratio. Cells were analyzed after 4 hr culture, n=183 cells across N=3 biological replicates. (G) Stiffness of gels measured using AFM. (H) SEM images of MD, MDx, LD and LDx gels, imaged at 15,000x and 30,000x magnification. Scale bar represents 2 µm for the left panel and 500 nm for the corresponding right panel. Statistical significance was determined using ANOVA, *p <0.05, **p < 0.01, and ****p < 0.0001.

Next, we cultured human foreskin fibroblasts (HFF) on each matrix condition to assess their response to matrix degradation. Fibroblasts were chosen because they are early responders to ECM remodeling^[26–28]^. HFF spreading was assessed after 4hr to capture the early response of cells to degradation. Cell spread area, circularity and aspect ratio were used for characterizing the mechanosensation of the cells. Cells on MD gels exhibited an average spreading area of 1741 µm^2^ and elongated morphologies, while cells on the LD gels exhibited rounded morphologies with a lower average spread area of 1007 µm^2^ (Figure 1C-F). After collagenase treatment, cells on MDx gels showed less spreading and more round morphologies compared to cells on the MD gels (Figure 1C-F). However, cells on LDx gels exhibited similar spreading, circularity, and aspect ratios to LD gels. These results suggest that cells sense changes in the matrix mediated by degradation and that these changes may correspond to mechanical properties.

To investigate this, we first measured the stiffness of each gel condition using AFM. Since cells are thought to apply nano newton forces on ECM^[29,30]^, we used a micron sized indenter and applied nanonewton forces to measure cell scale effects (see Methods). MD gels were slightly stiffer than the LD gels, 62 ± 13 Pa versus 38 ± 18 Pa, respectively (Figure 1G). We wondered if this observation could be due to the presence of thicker fibrils in MD gels, increasing the fibril bending stiffness. Indeed, scanning electron microscopy (SEM) of MD gels revealed relatively thicker fibril widths (0.13 ± 0.02 µm) when compared to the LD gels (0.097 ± 0.01 µm, Figure 1H and S1). After degradation, MD gels showed a reduction in stiffness from 62 ± 13 Pa to 30 ± 8.5 Pa, but LD gels did not exhibit changes in stiffness after degradation (35 ± 21 Pa, Figure 1D).

SEM revealed that collagenase treatment of MD gels significantly reduced fibril widths (MDx with 0.09 ± 0.008 µm, Figure 1H). However, collagenase treatment of LD gels did not change fibril widths (LDx) (Figure S1). These data show that treatment of MD gels with collagenase (MDx) resulted in an average stiffness and fibril width that was not significantly different from LD and LDx gels, which didn’t change with collagenase treatment. Cells spread less on the lower stiffness thinner fibril conditions (MDx, LD, LDx).

To decouple the influence of stiffness from degradation, we designed a control condition that exhibits the same stiffness as MDx gels but is not treated with collagenase. For this, we used 1.5 mg/mL collagen to produce gels and subsequently treated them with 0.01% glutaraldehyde for 5 minutes. The resulting gels (MDx-Control) exhibited the same stiffness as the MDx gels (Figure 1G), yet HFFs exhibited more spreading and elongated morphologies on MDx-Control gels than MDx gels (Figure 1C-F). The response of the cells to MDx-Control gels was similar to their behavior on MD gels, despite the MDx-Control gels exhibiting half the stiffness of MD gels. This suggested that although the stiffness of collagen gels reduces slightly after degradation, this is not the main cue driving the cellular mechanotransduction response to degradation, at least in the low stiffness ranges used here (30-60 Pa). This led us to hypothesize that cells may be sensing differences in stress relaxation post-degradation.

### 2.2 Degradation mediates enhanced stress relaxation in native collagen gels

AFM measurements showed that MDx gels exhibited greater stress relaxation compared to MD gels, while MDx-Control gels showed similar relaxation to the MD gels (Figure 2A). On the other hand, LD gel relaxation behavior was similar to LDx gels and MDx gels (Figure 2A). These data show that changes in stress relaxation correlate with matrix degradation. Moreover, enhanced stress relaxation (LD, LDx, MDx) correlates with less cell spreading (Figure 1D-F). To further quantify the stress relaxing behavior of the gels, we fitted the standard linear solid (SLS) model to the relaxation curves (Figure S2). The SLS model characterizes relaxation using equilibrium force (long-term) and relaxation time ^[32]^. Between the MD and LD gels, we observed no difference in the equilibrium force (F∞) (Figure 2B). However, the relaxation time τ showed an increase for LD indicating that these gels are slightly more viscous in nature (Figure 2C). For the MDx gels compared to their non-degraded counterpart (MD), an obvious increase in τ and a decrease in F∞ was seen. Specifically, the average F∞ and τ for MDx gels were 0.54 and 2 seconds, respectively, compared to 0.7 and 1.4 seconds for the MD gels (Figure 2B, 2C). LDx gels compared to LD gels did not show any difference in either of the SLS parameters. Similarly, the MDx-Control gels F∞ and τ were comparable to the MD gels. Overall, the SLS model reveals an increase in both F∞ and τ for the MD gels after degradation, which are similar to the F∞ and τ values of LD and LDx gels, and may explain the observation of less cell spreading MDx, LD, and LDx gels.

**Figure 2.**
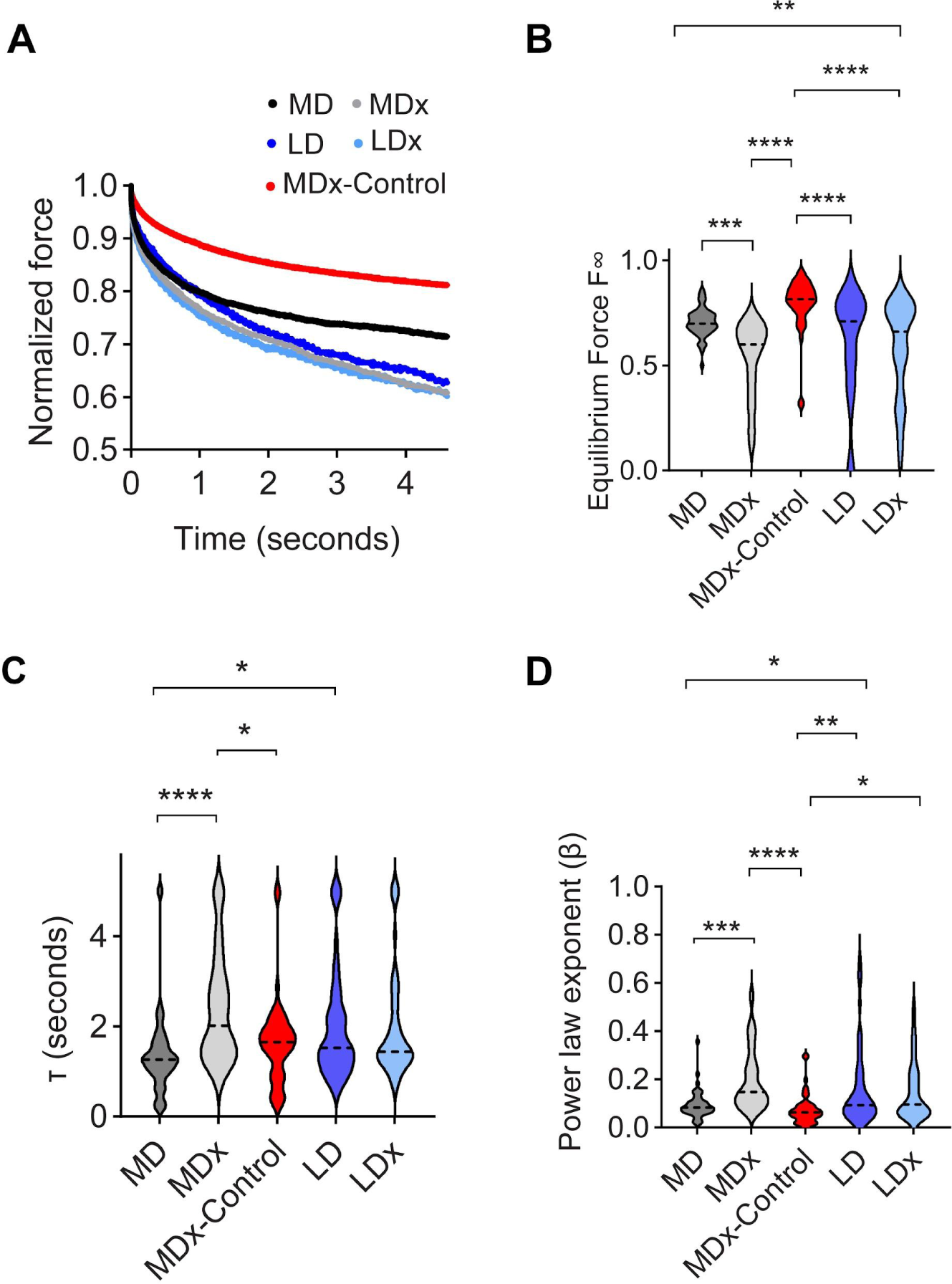
Differences in degradability of collagen gels correspond to differences in stress relaxation. MD - More degradable, MDx - More degradable gels with collagenase treatment, LD - Less degradable, LDx - Less degradable with collagenase treatment, MDx-Control - same stiffness as that of MDx gels. (A) Average of relaxation curves. (B) Standard linear solid parameter F_∞._ (C) Parameter Τ relaxation time. (D) Power law rheology fit parameter β. The data are obtained from n=3 independent samples, N = 15 measurements per sample. For all graphs, statistical significance was determined by ANOVA, *P <0.05, **P < 0.01, ***P < 0.001, and ****P < 0.0001.

The SLS model failed to capture the initial time points (t < 0.1 s, Figure S2), which is consistent with previous experiments on cells^[33]^. Therefore, we also performed a power law rheology (PLR) model fit (Figure 2D) to the relaxation curves. This yielded better fits than SLS (Figure S3). Higher values of the beta exponent in PLR corresponds to enhanced relaxation and more fluid-like behavior. The beta exponent showed an increase from an average of 0.09 to 0.2 for MDx gels compared to MD gels, suggesting a more fluid-like behavior post-degradation. Similar to the SLS fits, PLR fits showed that LD and LDx gels did not exhibit differences in their beta exponents. Together, these experiments show that the degradability of collagen gels is translated into significant changes in stress relaxation after collagenase treatment.

### 2.3 Design of hydrogels with varying degradability results in tunable stress relaxation post collagenase exposure

Next, we sought to rationally design synthetic hydrogels such that degradability tunes stress relaxation to mimic the effects we observed in our collagen model system. For this, poly (vinyl alcohol) (PVA) (molecular weight in range 40-300 kDa per manufacturer) gels of tunable degradability were prepared by varying the ratio of MMP degradable to non-degradable cross-links. We prepared two gel formulations denoted as collagenase degradable (CD) and non-degradable (ND) gels. To distinguish the influence of stiffness, we aimed to create the CD hydrogel formulation that exhibits similar stiffness as the ND gels but shows difference only in relaxation post degradation. The CD gels comprised 75% of MMP-degradable cross-links (Pro-Leu-Gly-Leu-Trp-Ala) and 25% of non-degradable PEG cross-links. The ND gels were made with 100% non-degradable PEG cross-links (Figure 3A). The stiffnesses of CD and ND gel formulations, around 3.5 kPa (Figure 3B), were not significantly different.

**Figure 3.**
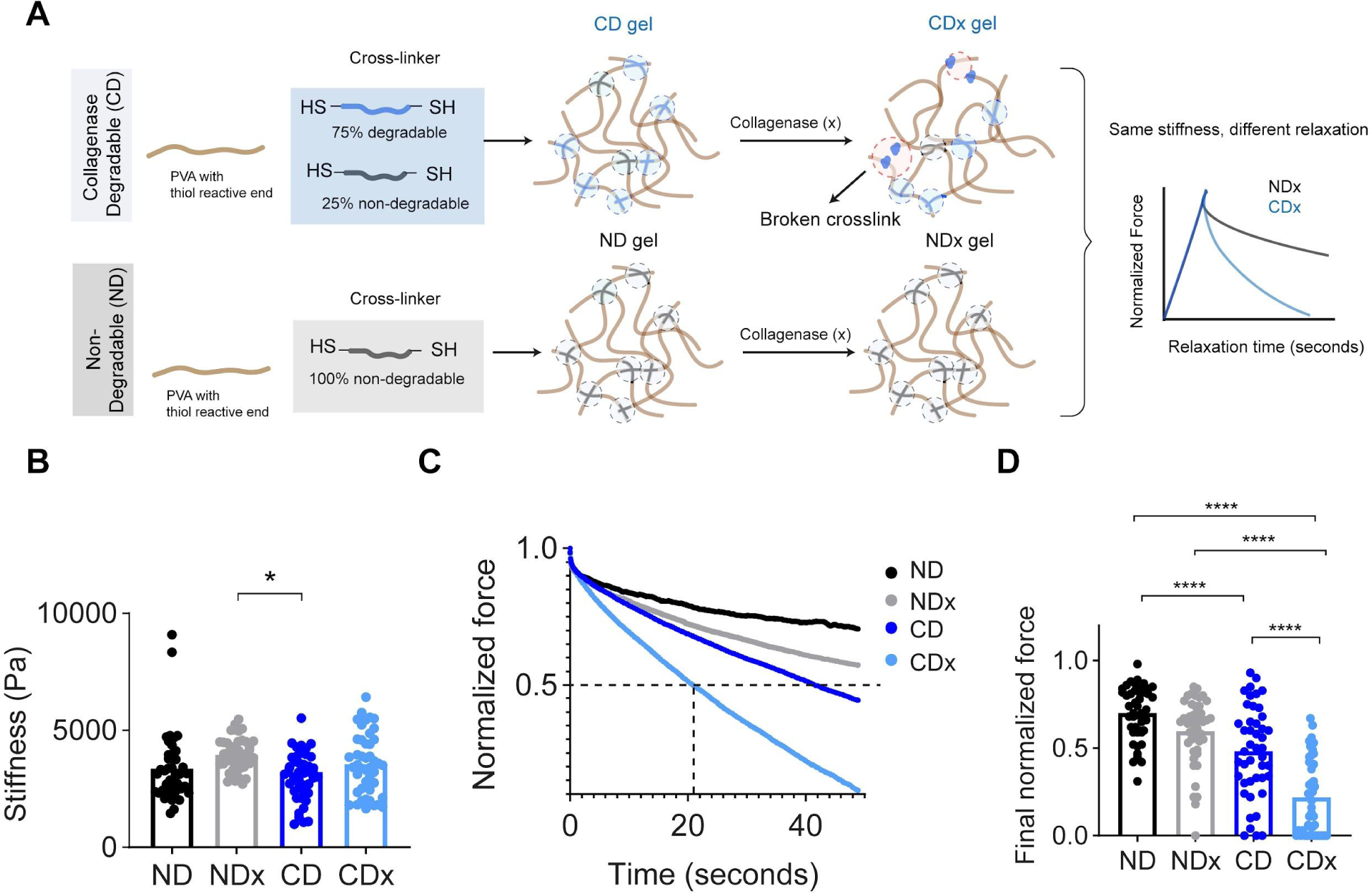
Rational design of synthetic stress relaxing gels as a function of degradability. (A) PVA gels of tunable degradability are generated by varying the ratio of non-degradable to degradable cross-links (created with BioRender.com). (B) Stiffness of hydrogels, N=45 across three independent samples, ND - non-degradable, NDx - non-degradable treated with collagenase, CD - collagenase degradable, and CDx - collagenase degradable treated with collagenase. (C) Stress relaxation of hydrogels assessed using AFM. (D) Final normalized force, n=15 measurements per gel. For all graphs, statistical significance was determined using ANOVA, *P <0.05 and ****P < 0.0001.

The gels were then subjected to 5 min of collagenase exposure at 37°C followed by assessment of mechanical properties. As a control, we confirmed that gels with 100% MMP-degradable cross-links exhibited a reduction in stiffness indicating degradation (Figure S4). However, the combination of non-degradable and degradable cross-links used for the CD gel did not exhibit significant differences in stiffness after degradation (CDx). Similarly ND gels did not exhibit differences in stiffness post treatment with collagenase (NDx) (Figure 3B).

Next, we characterized relaxation (Figure 3C) and quantified the time taken to relax to half of the maximum force (t_1/2_)^[34,35]^. For the cases of ND and NDx, >90% and 80%, respectively, of the relaxation data did not reach half of its maximum value. In contrast for CD and CDx gels, the percentages were closer to 70% and 10% respectively (Figure S5). For NDx and CD gels, the time to reach half of the maximum value was mostly greater than 40 seconds, indicating that the gels are predominantly slow-relaxing ( Figure S5). We also compared the relaxation amplitudes at the end of the experiment, which showed an average value of 0.7 for ND and around 0.5 for both NDx and CD gels (Figure 3D). The raw relaxation curves further indicated that CDx gels exhibited relatively homogeneous relaxation dynamics when compared to NDx, CD and ND gels (Figure S5). This shows that the enzyme treatment results in spatially uniform mechanical properties. Figure S6 shows that the relaxation trend of CDx gels is distinct from the other conditions and can be observed within 5 seconds of force application. We fitted a PLR model to the first 5 seconds of data and we found that the power law exponent increases for CDx gels indicating viscous behavior. Thus, after enzymatic degradation, our synthetic degradable gels show faster and enhanced relaxation while our non-degradable gels relax slower.

### 2.4 Fibroblasts exhibit less focal adhesions on degradable substrates after collagenase treatment

Since HFFs responded to degradation mediated stress relaxation of collagen gels, we speculated that they should respond in a similar way on our synthetic gels. To enable cell attachment, we conjugated 0.5 mM thiolated RGD to PVA before cross-linking. RGD modified hydrogels did not show differences in stiffness and relaxation compared to their unmodified counterparts (Figure S7).

First, we cultured HFFs on CD and ND gels. As expected, the cell area, circularity and aspect ratios showed no significance between the two conditions after 4 hr (Figure S8). Then, we proceeded to evaluate cell spreading on collagenase treated conditions. HFFs exhibited larger cell spread areas with an average of 2547 µm^2^ on non-degradable slow-relaxing NDx gels when compared to smaller spread areas (average -1336 µm^2^) on fast-relaxing CDx gels. Moreover, HFFs were more circular on CDx gels when compared to the NDx gels. Focal adhesions (FA) were then characterized using a vinculin antibody stain (Figure 4F,G). On NDx gels, focal adhesions were generally punctate, characteristic of mature focal adhesions^[36]^. The average focal adhesion (FA) number per cell was 59 and the mean area occupied by FA per cell was around 3.1%. In comparison, the FA on CDx gels were diffuse, suggesting less focal adhesion formation. Quantification of FA on CDx gels showed an average FA number per cell was 13 and average area occupied by FA was 1.7% indicating cells were largely less adhered. Overall, fibroblasts sense changes in stress relaxation on degraded CDx gels by spreading less and forming less FA.

**Figure 4.**
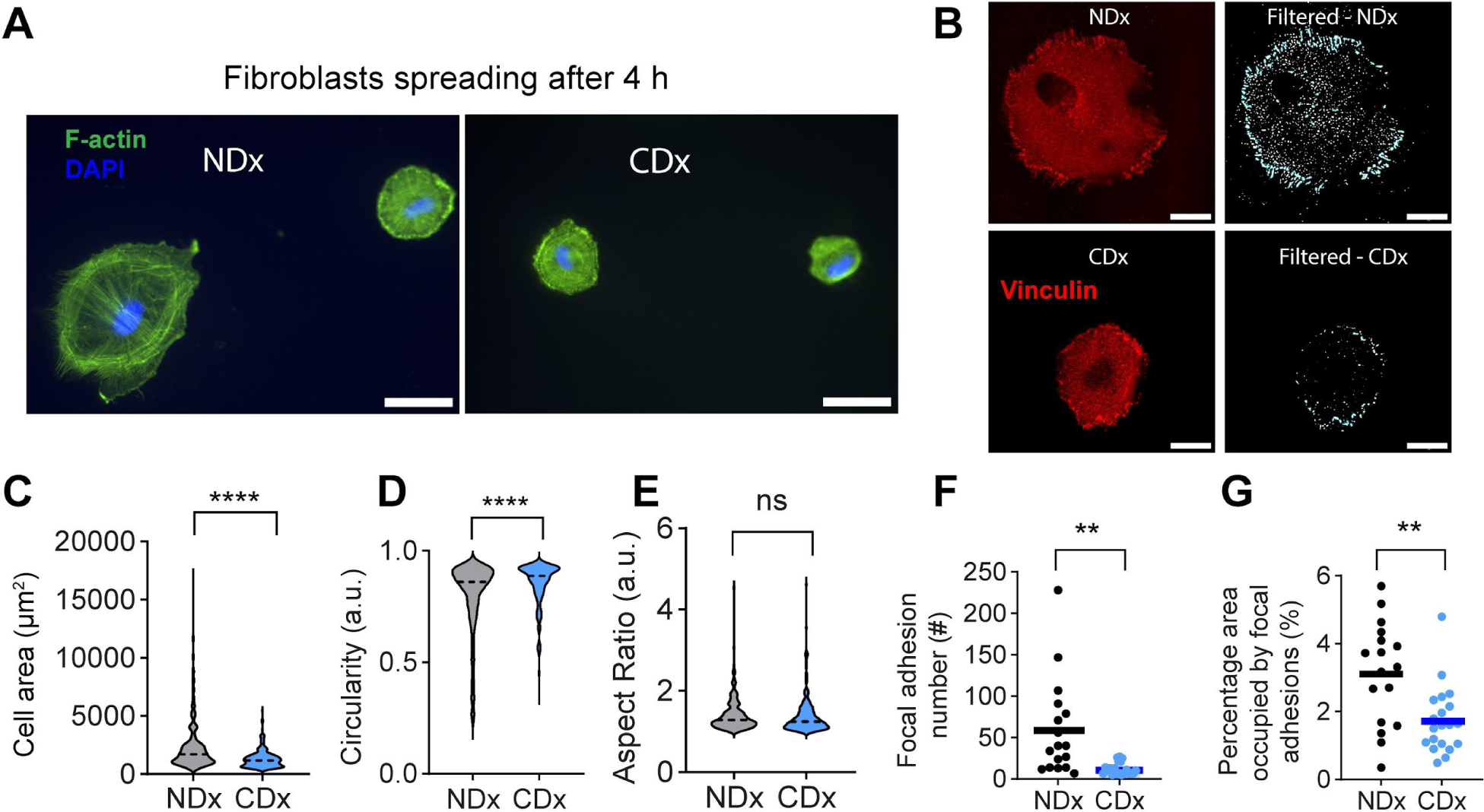
The fibroblast response to degradation of synthetic degradable gels is similar to their response to degradation of collagen gels. NDx - Non-degradable gels with collagenase treatment, CDx - Collagenase degradable gels with collagenase treatment. Human foreskin fibroblasts (HFF) spreading on gels (A) Phalloidin (green) and DAPI (blue) stained images. Scale bar = 50 µm. (B) Vinculin focal adhesions and corresponding filtered images showing adhesions. Scale bar = 20 µm. Quantification of cell spreading: (C) Area, (D) Circularity, and (E) Aspect ratio, n=247 cells measured across N=3 biological replicates. (F) Number of focal adhesions per cell with area between 1-10 µm^2^ and (G) Percentage of area occupied by focal adhesions per cell quantified using vinculin staining, n=17 and n=21 cells analyzed for NDx and CDx conditions across N=3 biological replicates. For all graphs, statistical significance was determined using student’s t-test, **P < 0.01 and ****P < 0.0001.

### 2.5 Cell response to stress relaxation induced by degradation is cell type dependent

Since the impact of stress relaxation has been shown to elicit differential responses depending on cell type^[37]^, we next asked if different cell types were capable of sensing stress relaxation induced by degradation. Previous studies investigating cell responses to substrate viscoelasticity have typically used a 24 hr time point, especially for adherent cell types ^[38–41]^. So, for comparison, we chose to assess cell spreading of two adherent cell types: C2C12 myoblasts and mammary epithelial cells (MCF10A) after 24 hr culture.

First, we cultured C2C12 myoblasts on NDx and CDx gels. CDx gels inhibited cell spreading (area - 1090 ± 650 µm^2^) compared to NDx gels (area- 1486 ± 877 µm^2^) (Figure 5A-D). Similarly, the cells were less elongated with an average aspect ratio of 2 on CDx gels compared to NDx gels (average aspect ratio - 2.9). In summary, myoblasts responded to changes in stress relaxation accompanying degradation in a similar way as fibroblasts.

**Figure 5.**
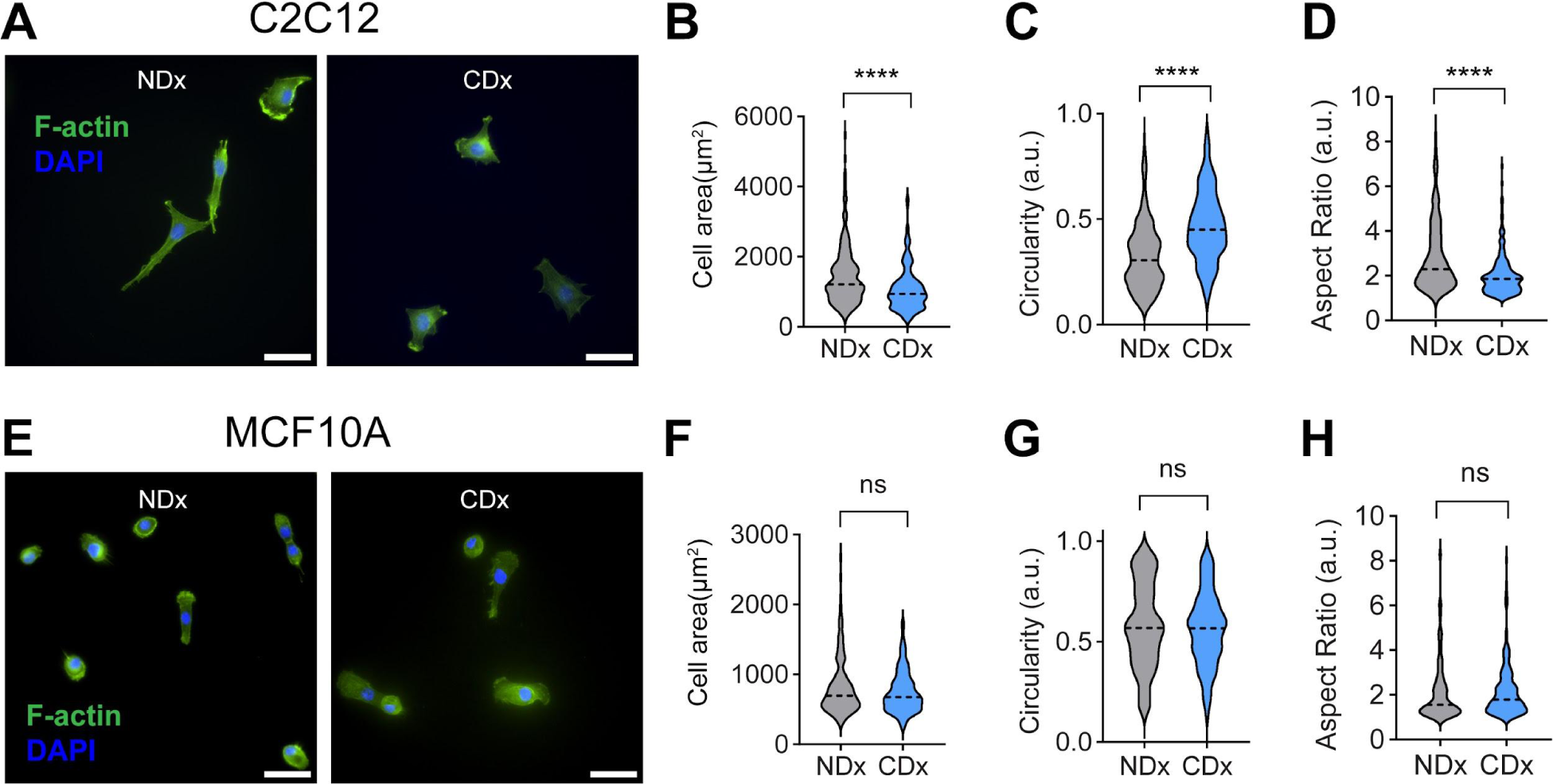
Degradation induced stress relaxation effects on cell spreading are cell type dependent. (A) C2C12 myoblasts spreading on NDx and CDx gels. Assessment of C2C12 spreading: (B) Area, (C) Circularity, and (D) Aspect ratio after 24 hr of culture, n=193 cells across N=3 biological replicates.(E) MCF10A spreading on NDx and CDx gels. Quantification of MCF10A spreading: (F) Area, (G) Circularity, and (H) Aspect ratio after 24 hr culture, n=143 cells across N=3 biological replicates. For all graphs, student’s t-test was performed, ****P < 0.0001.

Previous studies show that mammary epithelial cells (MCF10A) exhibit round morphologies on less than 1 kPa substrates and elongated morphologies on greater than 1 kPa substrates ^[42]^. Since the gel stiffness of NDx gels were around 3.5 kPa, most cells exhibited spread and elongated morphologies (Figure 5E), as expected. Interestingly, MCF10As on both CDx and NDx gels exhibited similar cell spread areas, circularity and aspect ratios (Figure 5F-H) indicating that MCF10A did not respond to substrate viscoelasticity induced by degradation.

## 3 Discussion and conclusion

Both synthetic and native ECMs are often generated with the goal of presenting precise elastic and viscoelastic mechanical cues to cells. Our study shows that when such systems are degradable, degradation can alter these specified properties. Using AFM, we characterized the cell scale-relevant changes in mechanical properties resulting from degradation and showed that in both synthetic and collagen matrices, stress relaxation was increased. This change was readily sensed by cells. Fibroblasts and myoblasts readily responded to degradation-induced changes in stress relaxation by spreading less and forming fewer focal adhesions on gels; however, mammary epithelial cells did not. Cell response to substrate viscoelasticity is known to depend on cell type and the dynamics of relaxation. For instance, Mandal et al^[37]^, utilized stress relaxing polyacrylamide gels and assessed the spreading behavior of normal hepatocytes and hepatocellular carcinoma cells. While normal hepatocytes showed less spreading on stress relaxing substrates, carcinoma cells showed enhanced spreading on stress relaxing substrates when compared to elastic substrate counterparts of the same stiffness.

Similar to our observations, a previous study of 3T3 fibroblasts cultured on polyacrylamide gels of varying viscous dissipation showed less spreading on substrates that exhibited enhanced relaxation^[44]^. On the other hand, fibroblasts have also been shown to spread better on stress relaxing alginate substrates compared to their elastic counterparts, depending on ligand concentration^[27]^. A previous study of C2C12 myoblasts on alginate matrices also showed that enhanced stress relaxation promotes cell spreading and elongated morphologies^[39]^, which runs counter to our observations. However, in the prior study, C2C12 cells were cultured on elastic alginate gels (little relaxation) and slow stress-relaxing gels ( t_1/2_ > 120 seconds). In comparison, our NDx gels and CDx gels exhibited an average t_1/2_ of 43 and 23.5 seconds respectively. So, the slow-relaxing matrices from Bauer et al^[39]^ are similar to our NDx gels, which did promote more cell spreading and higher aspect ratios. For MCF10A mammary epithelial cells, our results indicated that they did not respond to our fast relaxing CDx substrates (t_1/2_ of 23.5 seconds) induced by degradation in 2D at a 24 hr time point. A recent study which used mammary epithelial cells cultured on poly(acrylamide) gels of 2 kPa stiffness with varying viscous dissipation showed cell spreading was inhibited on highly dissipating matrices corroborating our findings^[41]^. However, other research showed that MCF10A spheroids exhibited enhanced spreading in 3D fast relaxing substrates (t_1/2_ of 30 seconds) compared to slow relaxing substrates (t_1/2_ of 350 seconds)^[45]^. The inconsistency of these results may be attributed to differences in ligand density, which has been suggested to modulate the viscoelastic contribution to spreading^[27,39]^. Another important difference to consider is the characterization technique, i.e. bulk (compression or rheology) vs. local AFM and the method of force application (shear vs. compression), which makes it difficult to compare between the relaxation timescales. We argue that cell scale relevant characterizations are needed to gain a deeper understanding of timescales that are physiologically relevant. In summary, cellular responses to stress relaxation vary, even for cells of the same type. These responses appear to depend on relaxation dynamics, which may differ based on how relaxation is assessed, as well as other factors like ligand density.

While enzymatic degradation of ECM can lead to a reduction in stiffness^[43]^, we found that it can lead to altered stress relaxation independent of a reduction in stiffness. The extent of each type of mechanical change is likely to depend on the concentration and time of exposure to the degrading enzyme as well as its activity and potentially its mechanism of degradation. Together, this suggests that the matrix mechanics felt by cells in native and synthetic degradable ECMs significantly depends on the susceptibility of the matrix to degradation and the capability of the cells to produce degrading enzymes. This also means that degradability is a feature that can be designed to present cell and MMP-specific stress relaxation responses over time. In our synthetic system, changes in relaxation post degradation were observed as early as 5 seconds after force application. The gels became fast relaxing after degradation. Tuning the architecture of the degradable crosslinker can potentially offer additional control over specific stress relaxation times.

To tune collagen gel degradability in our study, we used a previously developed approach that employs macromolecular crowding. It is interesting to note that degradability correlates with macromolecular crowding-induced changes in collagen architecture. Gels that are more degradable had thicker fibers and showed more change in fibril widths after degradation compared to less degradable gels. While deciphering the mechanisms underlying the difference in degradability and fibril network mechanics is beyond the scope of this work, we speculate that the thickness of collagen fibrils may play a major role in modulating the degradability and the subsequent viscoelasticity of collagen matrices. It would be interesting to investigate whether this characteristic of collagen architecture may potentially explain differences that others have observed in the relaxation behavior of advanced glycan end products (AGE)-crosslinked collagen matrices^[46]^. In the future, it would also be interesting to investigate if these relationships hold true for other fibrous ECM matrices such as fibrin.

In conclusion, this work demonstrates that even light exposure of hydrogels to degrading enzymes, which doesn’t alter stiffness, can have a significant impact on their stress relaxation behavior. Accordingly, degradability was leveraged as an engineering design variable to tune stress relaxation and cellular mechanotransduction. This holds significant implications for the design of future biomaterials. For example, the degradability of hydrogels could be engineered in cell type or MMP-specific ways to spatially and temporally control stress relaxation properties. This work also demonstrates that cellular interactions with materials through local degradation could change the material to become biophysically different from its initial characterization. Since material properties are typically characterized after manufacture without accounting for cellular modification, adjusting for degradation effects could be crucial for engineering future biomaterials in order to maintain a desired environmental condition to control cell behavior. This may be especially important in fields like regenerative medicine, where degradable scaffolds are used to promote cell infiltration and repair injured tissues.

## 4 Experimental section

### Chemicals

3-D Life PVA-PEG kit hydrogel kit SG (Cat. No.:G82-1), 3-D Life PVA-CD kit hydrogel kit SG (Cat. No.:G83-1) and 3-D Life RGD peptide (Cat.No.: 09-P-001) were purchased from Cellendes GmbH.

High concentration, rat tail acid extracted type I collagen was procured from Corning (Corning, NY). PEG (8000 Da) was ordered in powder form from Sigma-Aldrich (St Louis, MO) and reconstituted in PBS (Life Technologies, Carlsbad, CA) immediately before usage with a final concentration of 100 mg/mL.1× reconstitution buffer composed of sodium bicarbonate, HEPES free acid, and nanopure water.

### Preparation of collagen hydrogels with different architectures

First, PEG of required amounts to make a 2 or 8 mg/mL final concentration (denoted as P2 or P8) was added to DMEM. This is followed by addition of the reconstitution buffer and mixing well. Thereafter, collagen stock was added to the mixture to produce a final concentration of 2.5 mg/ml. Finally, pH of the final mixture was adjusted using 1 N NaOH,followed by incubation at 37 °C (∼45 minutes). Following polymerization, PEG was washed out of the gels by rinsing with DMEM 3× for 5 min each.

The gels were then degraded for 40 mins using 10 μg/mL of bacterial collagenase followed by rinsing with DMEM for 3x times.

### Preparation of synthetic hydrogels with tunable degradability

Gel consisting of 5 mM of thiol reactive PVA was formed using 3-D Life hydrogels according to the recipe shown in Table 1. For the gel preparation, the components listed in Table 1 are added in sequence. For cell culture, 0.5 mM of RGD is incubated with PVA for 20 mins at room temperature followed by the addition of the remaining components in Table 1 followed by mixing. After adding the cross-linker, the mixed solution was sandwiched between hydrophobic coverslip (DCDMS coated) and glutaraldehyde treated coverslip and allowed to react at 37°C for 45 minutes.

**Table 1.** Preparation of hydrogels with tunable degradability.

| Component | Non-degradable gel (ND) | Degradable gel (CD) |
| --- | --- | --- |
| Water | 15 µL | 15 µL |
| 10 x CB (7.2) | 3.75 µL | 3.75 µL |
| PVA | 7.5 µL | 7.5 µL |
| DMEM | 7.5 µL | 7.5 µL |
| CD-Link | - | 8.43 µL |
| PEG-Link | 11.25 µL | 2.82 µL |

The ND and CD gels were then treated with 10 μg/mL of Type I Clostridium histolyticum bacterial collagenase for 5 min and then washed with DMEM for 3x times followed by characterization.

### Atomic Force microscopy (AFM) on gels

AFM was used to characterize the local change in the mechanical properties of the hydrogels. AFM experiments were performed using a MFP-3D Bio atomic force microscope (Oxford Instruments) mounted on a Ti-U fluorescent inverted microscope (Nikon Instruments) using the force contact mode. Gold coated silicon nitride probes attached with a 12 μm borosilicate glass (Novascan, USA) of nominal spring constant 0.03 N/m was used for the measurements. The probes were calibrated using the thermal noise method in DMEM followed by indenting on a glass surface in DMEM for calibrating the deflection sensitivity using the built in Asylum Research Software (Igor Pro, Wavemetrics).

The gels were fabricated on the glutaraldehyde coated coverslips using the procedure outlined in a previous work ^[47]^ to ensure covalent attachment of gels during AFM indentations performed in liquid (DMEM). The thickness of the hydrogels produced were measured to be greater than 100 μm. For the stiffness measurements, the approach curve was fitted with a Hertz model for a spherical indenter geometry using the Asylum Research software which is described as follows^[48]^:

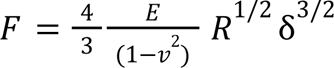

where F is the indentation force, R corresponds to the radius of the indenter, E is the Young’s modulus, and v is the Poisson’s ratio which is assumed to be 0.5 for elastic materials and δ is the indentation depth. The fits were performed on the first 35% of the force curve to quantify the Young’s modulus of the gels.

We sandwiched the pre-gel solution between the glutaraldehyde coated coverslip and a dichlorodimethylsilane (DCDMS) coated coverslip for the synthetic gels. For the collagen gels, the gels were casted onto glutaraldehyde coated coverslips without a hydrophobic coverslip cover. The thickness of the synthetic gels were around 150 µm for the synthetic gels and the collagen gels were around 400 µm. The diffusion coefficient of collagenase was approximately. 7.4x10^-7^ cm^2^/s^[49]^. The time to diffuse through 400 µm collagen gels was calculated to be around 36 minutes. Hence, we treated the collagen gels for 40 mins. On the other hand, the synthetic gels thickness was calculated to be around 130-150 µm. We calculated the time to diffuse through was around 5 mins. Hence, we used 5 mins for the synthetic gels.

For the force-relaxation experiments on synthetic gels, an indentation speed of 2 μm/s was used to achieve a relative trigger of 2 nN followed by holding the z-position of the probe for 50 seconds in which the force changes were captured before the probe was retracted. At least 15 viscoelastic measurements per sample were performed at random locations in the sample spaced at least 100 μm apart between locations. The viscoelastic measurements in which the approach curves were noisy were excluded for the analysis. For each condition, at least three individual gels were characterized. In NDx and CDx gels, some of the relaxation curves completely relaxed within 50 seconds of experimental acquisition time, and reached small negative values as the forces have completely relaxed. In such cases, the final normalized force value was taken as zero.

Due to the highly adhesive nature of collagen matrices causing rapid adhesion of the AFM tip to sample in the DMEM environment, we could not perform reliable measurements for more than 5 seconds for the collagen gels. Thus, we performed force-relaxation for a period of 5 seconds using the z-feedback loop of the AFM with a higher resolution (2 kHz).

### Relaxation curve analysis

The relaxation curves were analyzed by fitting either the SLS model or the PLR model using non-linear least squares fit. For the SLS model, the normalized force-relaxation curves were fitted using the following equation^[33]^:

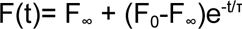

where F_∞_ refers to the final long-term equilibrium force, F_0_ refers to the instantaneous force and τ represents the relaxation time.

For the PLR model, the relaxation curves were fitted using the following equation^[33]^:

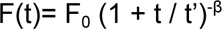

where F_0_ refers to the instantaneous force, t’ represents small time offset which does ont affect relaxation behavior, β represents the power law exponent.

### Scanning electron microscopy (SEM)

SEM was performed on the hydrogels using FEI SEM Apreo equipped with an ETD detector. Both the collagenase treated gels and the non treated gels were first fixed with 4% paraformaldehyde for 1 hr. This is followed by 3x rinsing in PBS for 10 min of each step of shaking. The gels were then rinsed twice with Milli-Q water for 15 min each. The fixed gels were then subjected to a series of dehydration steps in ethanol and hexamethyldisilazane (HMDS) using the protocol outlined in Raub et al^[50]^. Briefly, the gels were dehydrated first in ethanol dilution series: 30%, 50%, 70%, 90% and 100% for 15 min of each step. The hydrogels were then incubated in ethanol/HMDS dilution series: 33%, 50%, 66% and 100% for 15 min each. After the final incubation, the gels were allowed to dry in an aluminum foil for at least 1 day in the fume hood. The dried gel samples were then sputter coated with a Pelco SC-7 sputter coater with gold as the target. The gels were then imaged at 5 kV and 0.6 nA with 15,000x magnification. At least 12 images were acquired per sample, across three replicates for the quantification of mean width. For the width analysis, we used SIMpoly software^[50,51]^.

### Cell culture

Human foreskin fibroblasts (passages 11-24), C2C12 (passages 3-10), and MCF10A (passages 7-14) cell lines were purchased from ATCC and cultured in high glucose Dulbecco’s modified Eagle’s medium supplemented with 10% (v/v) fetal bovine serum (FBS, Corning, Corning, NY) and 0.1% gentamicin (Gibco Thermo Fisher, Waltham, MA) and maintained at 37°C and 5% CO_2_ in a humidified environment during culture and imaging. The cells were passaged every 2–3 days as required.

Cells were trypsinized and spun down and cultured on top of both synthetic and native collagen gels after washing 3x DMEM. For the HFFs, after 4 hr of culture, cells were fixed in 4% paraformaldehyde and then stained with DAPI and phalloidin for quantifying the cell area. For C2C12 and MCF10A cell lines, fixing and staining were performed after 24 hr.

### Focal adhesion staining and quantification

The focal adhesions were quantified using vinculin staining. For vinculin staining, fixed samples were blocked using 2% BSA for 2 h at room temperature and then incubated primary vinculin monoclonal antibody (1:250, V9131, Sigma) for 2 h. Following incubation, samples were rinsed 3x times in PBS to remove unbound primary antibodies. Then, samples were incubated with a secondary antibody conjugated with alexa-546 for 1 hr at room temperature followed by imaging after 3x times rinsing in PBS.

For the quantification of focal adhesions, we adopted a modified procedure from Tolbert et al. and Horzum et al. ^[52,53]^ Briefly, the images were background subtracted using rolling ball radius to 2 pixels in Fiji^[54]^. The local contrast of the image is enhanced by applying the CLAHE (Contrast limited adaptive histogram equalization) plugin with set parameters as: blocksize = 19, histogram bins = 256, maximum slope = 3. We then applied Gaussian blur with a sigma radius of 2 followed by a Mexican hat filter with sigma radius = 2. The filtered images were binarized using Otsu thresholding method. Finally, using the Analyze particles option we selected circularity from 0.00-0.99 and size above 1 µm^2^. The focal adhesion number reports size ranges 1-10 µm^2^ and the area per focal adhesion was calculated by dividing the total area of focal adhesions divided by the cell area obtained from the phalloidin staining.

### Cell area quantification

The cell area, circularity and aspect ratio were obtained using the images stained with phalloidin and DAPI. We used the Trackmate/Fiji plugin^[55]^ for all of our analysis.

## Author contributions

B.N.N. designed and performed all experiments and analyzed data. S.I.F conceived and supervised the project. S.I.F and B.N. wrote the manuscript.

We would like to thank Prof. Padmini Rangamani for the critical discussions that helped in shaping the manuscript. We would like to thank Prof. Adam Engler for the use of his atomic force microscopy instrument. We thank all the members of the Fraley lab for their critical feedback on this manuscript.

## Supporting information

**Figure S1.**
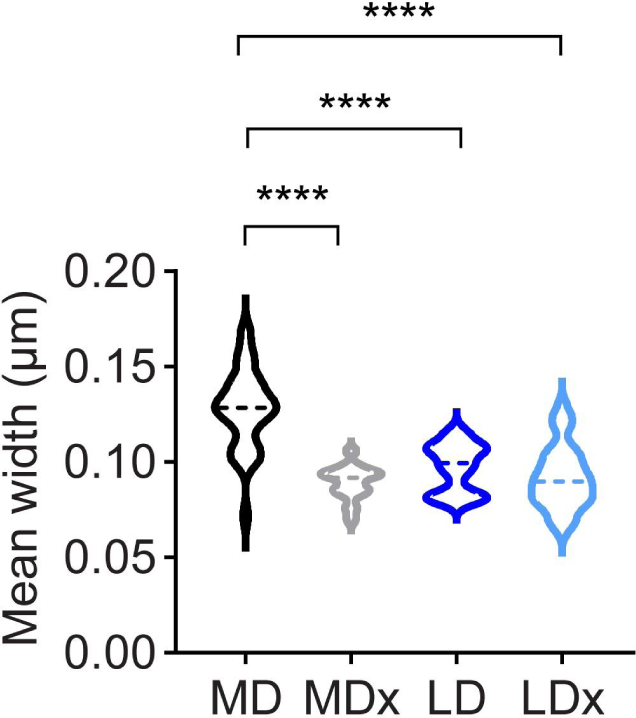
Mean widths of collagen fibrils assessed by scanning electron microscopy. The data show values obtained from at least 12 images obtained from three independent samples.

**Figure S2.**
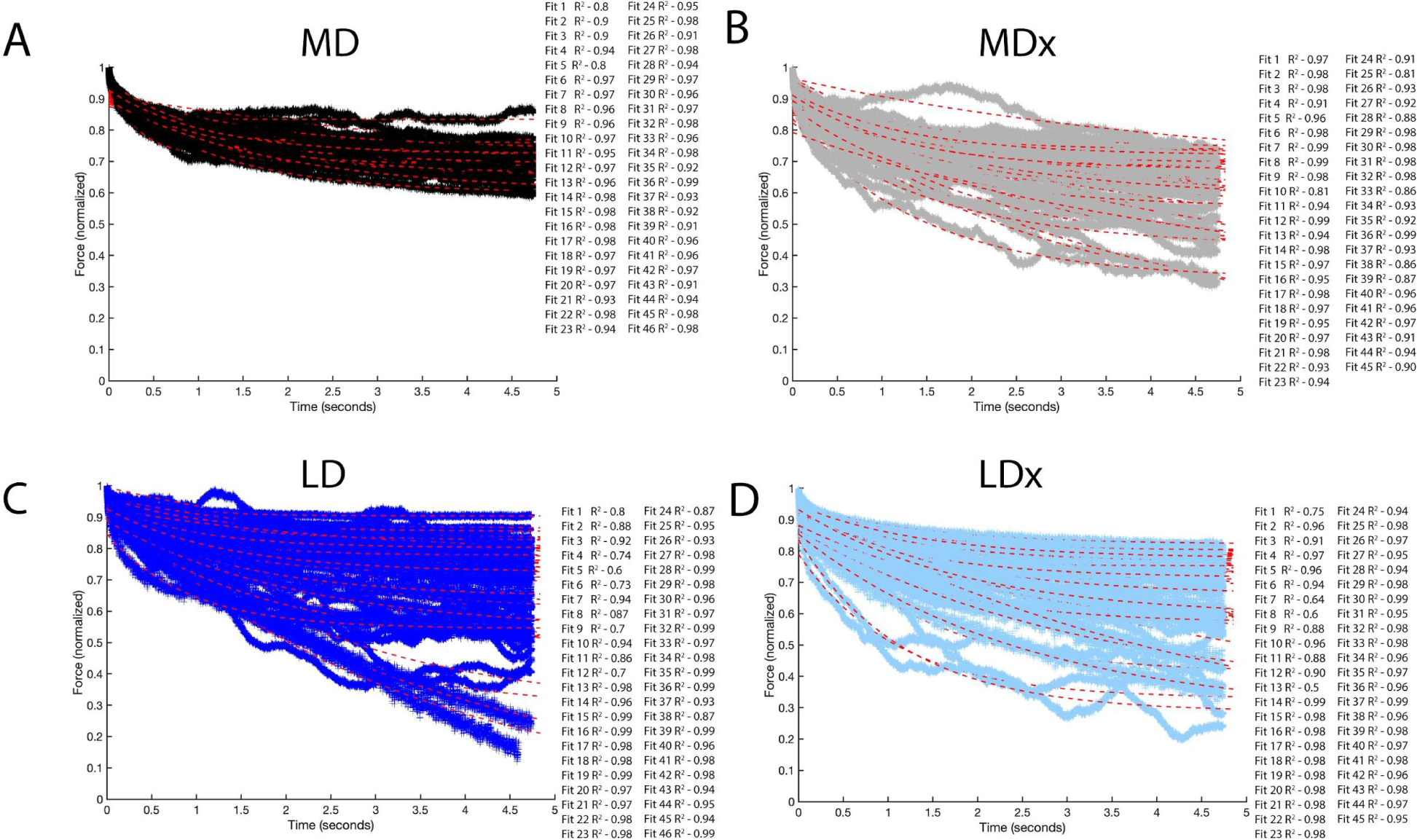
Standard linear solid fits for collagen gels.

**Figure S3.**
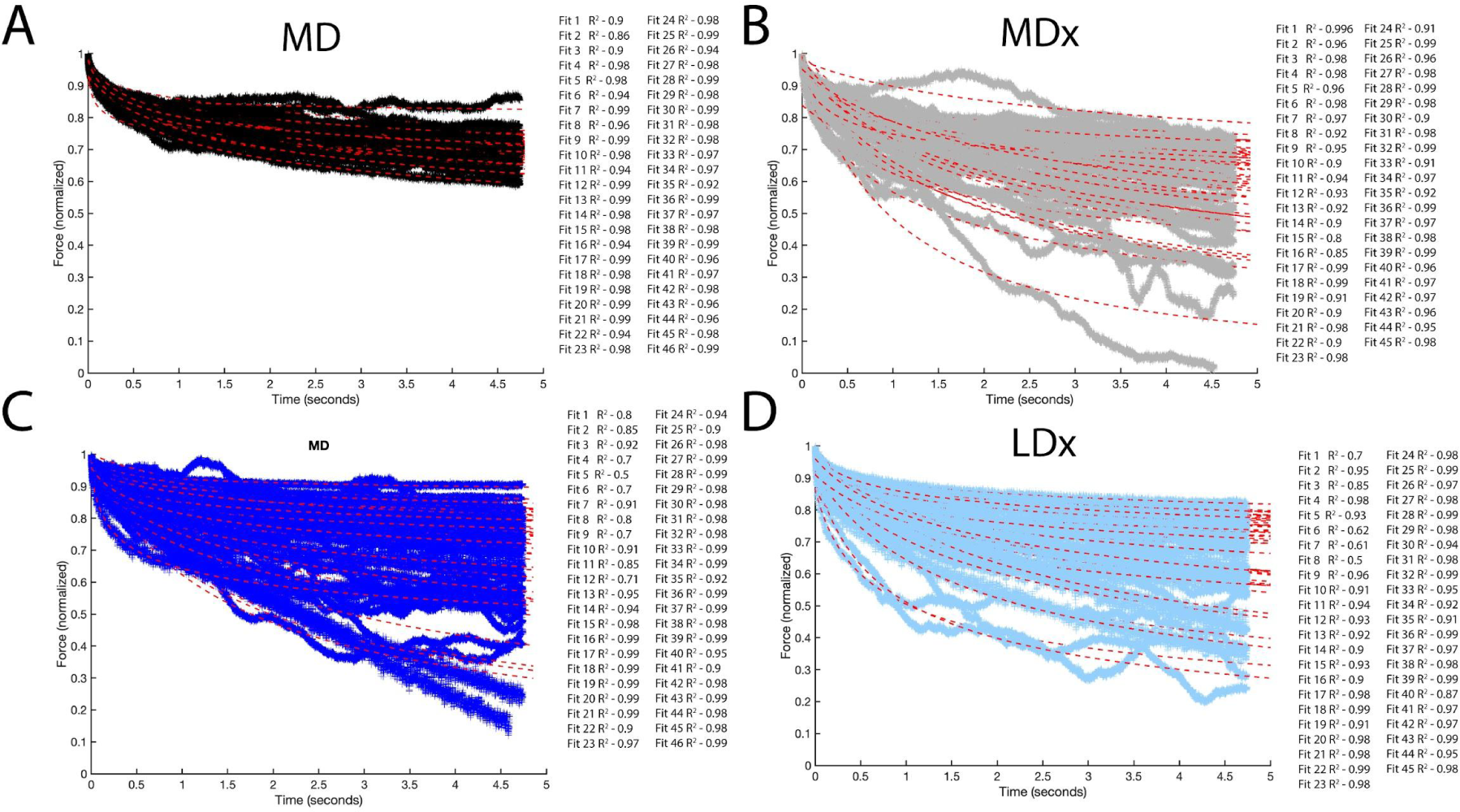
Power law fits for collagen gels.

**Figure S4.**
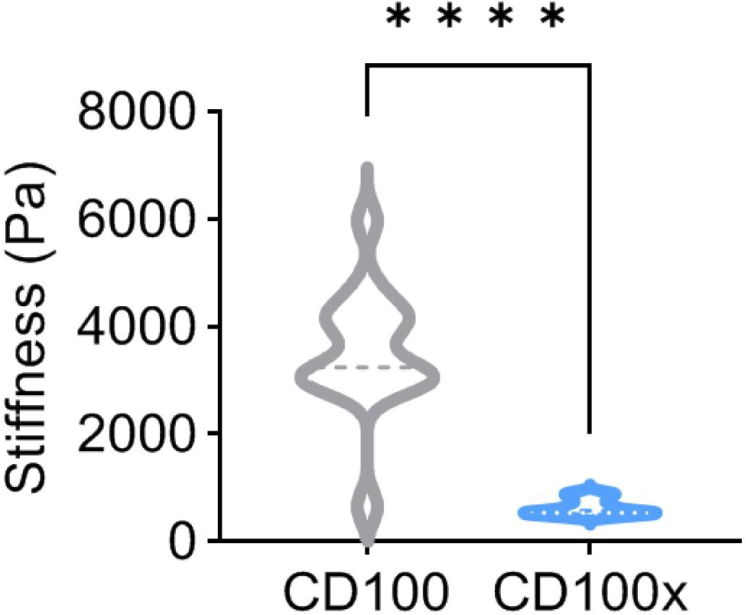
Stiffness of PVA gels made with 100% degradable cross-links. The gels exhibited significant reduction in stiffness after treatment with collagenase.

**Figure S5.**
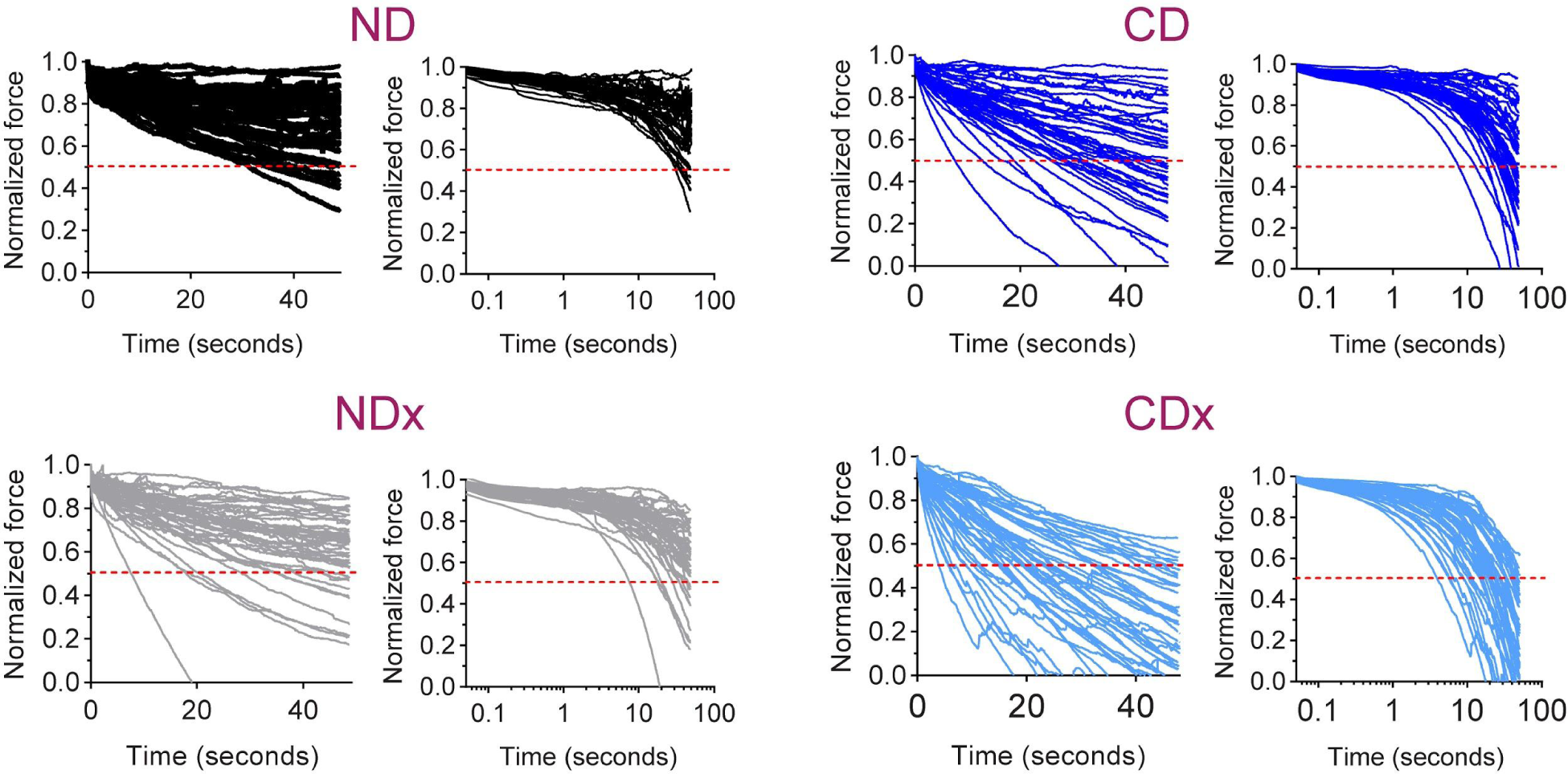
Raw relaxation curves of ND, NDx, CD and CDx with normalized force in linear scale and time axis in log-scale.

**Figure S6.**
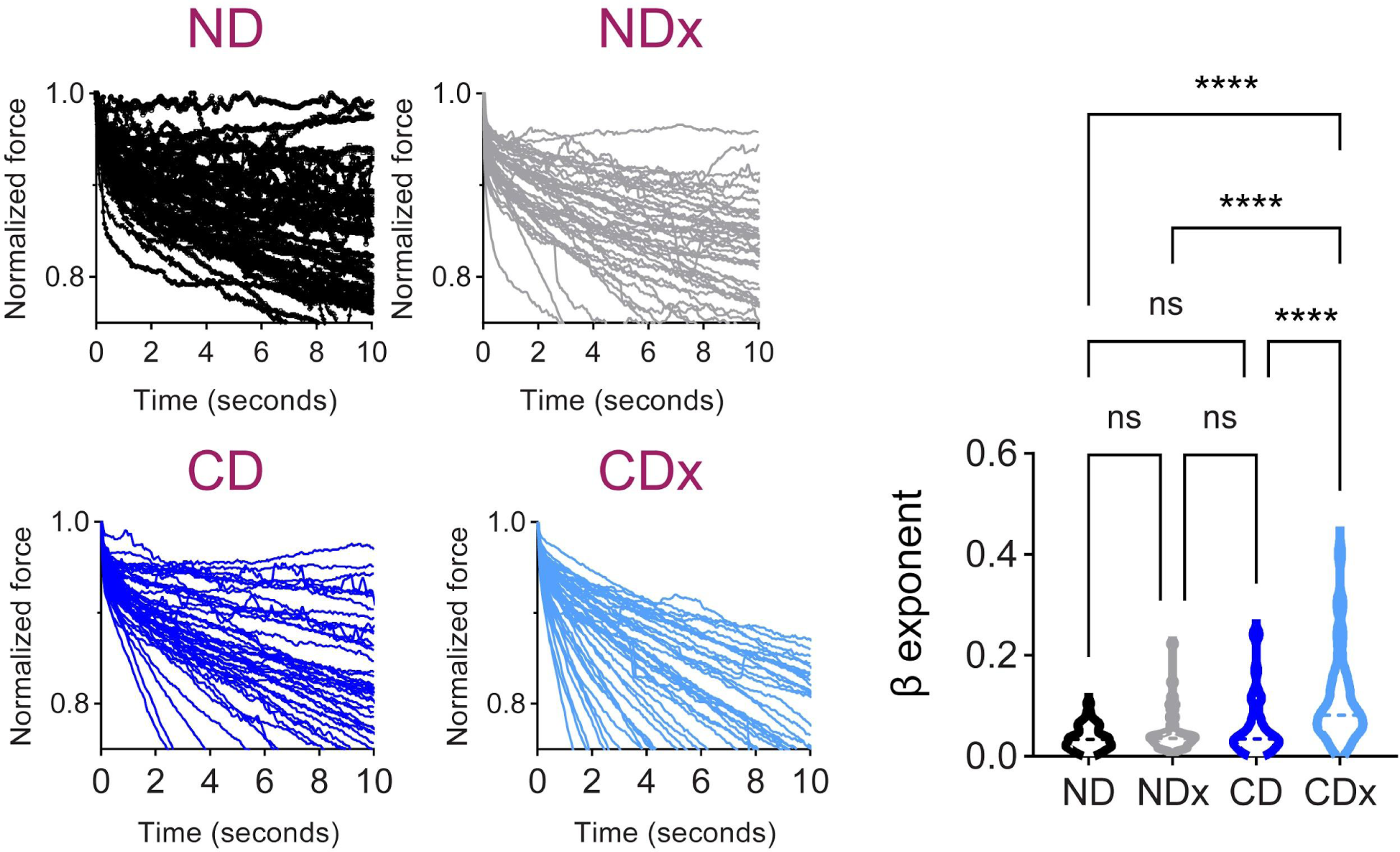
Power law rheology fits to the first 5 seconds of PVA gel data. One-way ANOVA analysis, ****P<0.0001.

**Figure S7.**
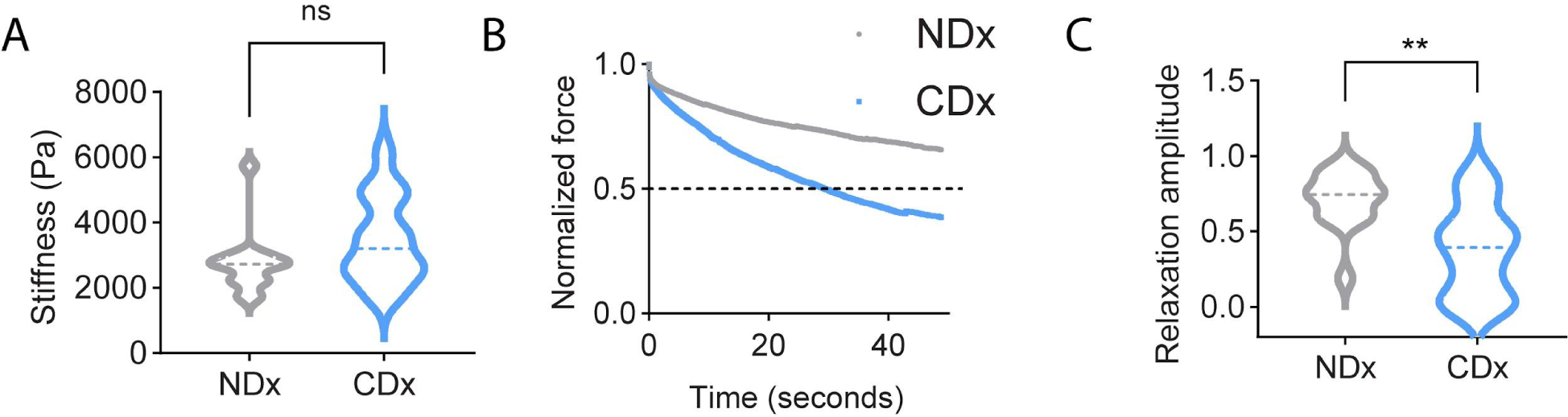
RGD modified hydrogels. (A) stiffness. (B) Average relaxation curve. The t_1/2_ was calculated to be around 25 ± 14 seconds. Three of the curves in CDx gel did not relax to more than half of the maximum value. None of the NDx gels reached 0.5 in the experiments. (C) Relaxation amplitude across data, **P < 0.01. N=15 measurements.

**Figure S8.**
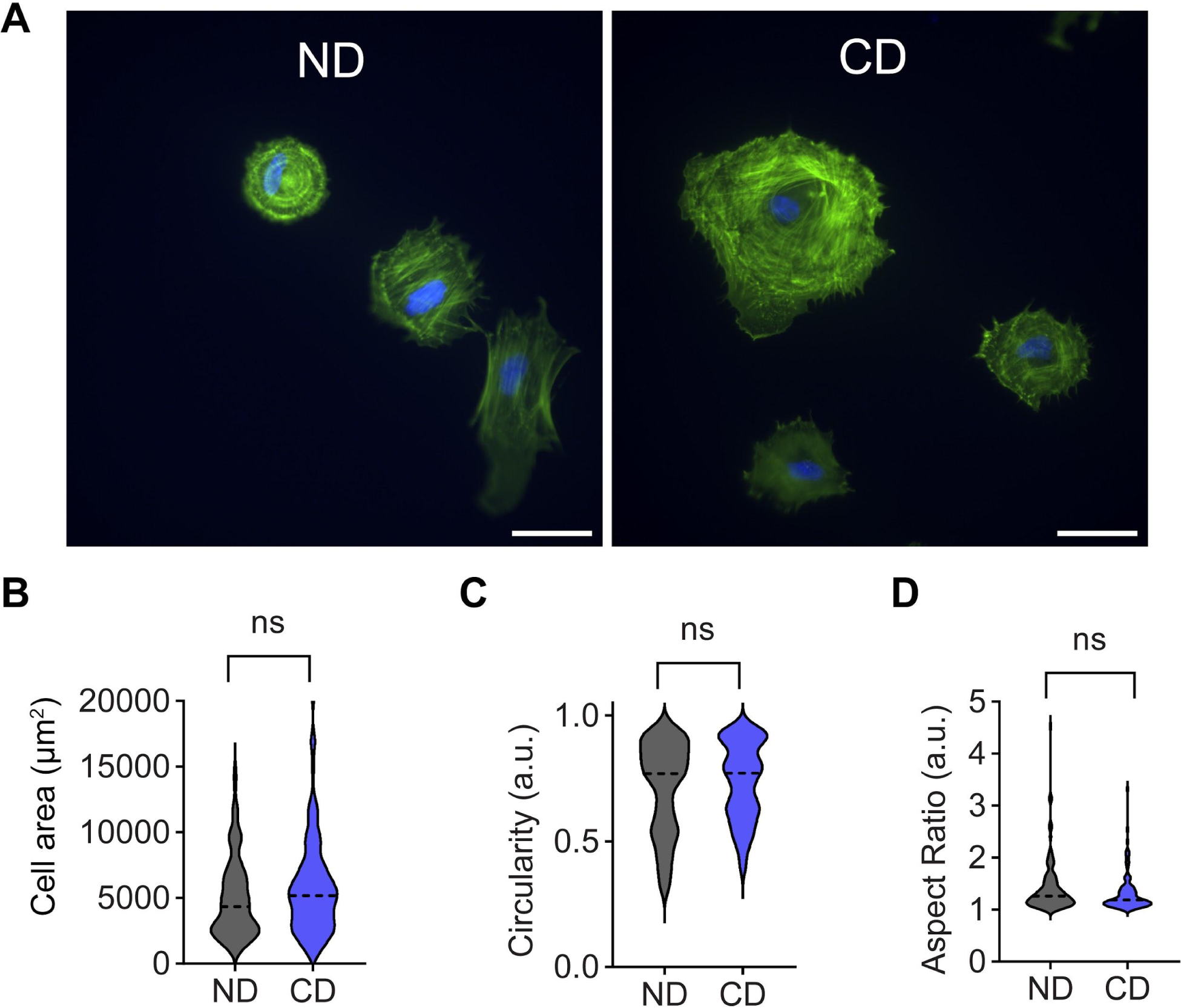
No difference in spreading was observed between Non-degradable (ND) and collagenase degradable (CD) gels. (A) HFF on ND and CD gels. Scale bar represents 50 µm. Cells spreading assessment: (B) Area, (C) Circularity, (D) Aspect ratio. n=93 cells, N=3 biological replicates. All graphs were analyzed using the student’s t-test.

## References

[1] M. Akhmanova, E. Osidak, S. Domogatsky, S. Rodin, A. Domogatskaya, Stem Cells Int. 2015, 2015, 167025.

[2] R. J. Pelham Jr, Y. l. Wang, Proc. Natl. Acad. Sci. U. S. A. 1997, 94, 13661.

[3] B. N. Mason, A. Starchenko, R. M. Williams, L. J. Bonassar, C. A. Reinhart-King, Acta Biomater. 2013, 9, 4635.

[4] E. M. Carvalho, S. Kumar, Acta Biomater. 2023, 163, 146.

[5] K. Adebowale, Z. Gong, J. C. Hou, K. M. Wisdom, D. Garbett, H.-P. Lee, S. Nam, T. Meyer, D. J. Odde, V. B. Shenoy, O. Chaudhuri, Nat. Mater. 2021, 20, 1290.

[6] A. G. Clark, A. Maitra, C. Jacques, M. Bergert, C. Pérez-González, A. Simon, L. Lederer, A. Diz-Muñoz, X. Trepat, R. Voituriez, D. M. Vignjevic, Nat. Mater. 2022, 21, 1200.

[7] L. Moretti, J. Stalfort, T. H. Barker, D. Abebayehu, J. Biol. Chem. 2022, 298, 101530.

[8] C. Loebel, R. L. Mauck, J. A. Burdick, Nat. Mater. 2019, 18, 883.

[9] A. Das, M. Monteiro, A. Barai, S. Kumar, S. Sen, Sci. Rep. 2017, 7, 14219.

[10] S. Khetan, M. Guvendiren, W. R. Legant, D. M. Cohen, C. S. Chen, J. A. Burdick, Nat. Mater. 2013, 12, 458.

[11] B. Trappmann, B. M. Baker, W. J. Polacheck, C. K. Choi, J. A. Burdick, C. S. Chen, Nat. Commun. 2017, 8, 371.

[12] S. K. Ranamukhaarachchi, R. N. Modi, A. Han, D. O. Velez, A. Kumar, A. J. Engler, S. I. Fraley, Biomater Sci 2019, 7, 618.

[13] J. Liu, H. Long, D. Zeuschner, A. F. B. Räder, W. J. Polacheck, H. Kessler, L. Sorokin, B. Trappmann, Nat. Commun. 2021, 12, 3402.

[14] S. Khetan, J. A. Burdick, Biomaterials 2010, 31, 8228.

[15] Y. Peng, Q.-J. Liu, T. He, K. Ye, X. Yao, J. Ding, Biomaterials 2018, 178, 467.

[16] O. Chaudhuri, L. Gu, D. Klumpers, M. Darnell, S. A. Bencherif, J. C. Weaver, N. Huebsch, H.-P. Lee, E. Lippens, G. N. Duda, D. J. Mooney, Nat. Mater. 2016, 15, 326.

[17] O. Chaudhuri, J. Cooper-White, P. A. Janmey, D. J. Mooney, V. B. Shenoy, Nature 2020, 584, 535.

[18] K. M. Schultz, K. A. Kyburz, K. S. Anseth, Proc. Natl. Acad. Sci. U. S. A. 2015, 112, E3757.

[19] B. Babaei, A. Davarian, S.-L. Lee, K. M. Pryse, W. B. McConnaughey, E. L. Elson, G. M. Genin, Acta Biomater. 2016, 37, 28.

[20] J. Winkler, A. Abisoye-Ogunniyan, K. J. Metcalf, Z. Werb, Nat. Commun. 2020, 11, 5120.

[21] K. E. Kadler, D. F. Holmes, J. A. Trotter, J. A. Chapman, Biochem. J 1996, 316 (Pt 1), 1.

[22] in Current Topics in Developmental Biology, Academic Press, 2018, pp. 107–142.

[23] S. R. Van Doren, Matrix Biol. 2015, 44-46, 224.

[24] H. Topol, H. Demirkoparan, T. J. Pence, Appl. Mech. Rev. 2021, 73, DOI 10.1115/1.4052752.

[25] I. Solomonov, E. Zehorai, D. Talmi-Frank, S. G. Wolf, A. Shainskaya, A. Zhuravlev, E. Kartvelishvily, R. Visse, Y. Levin, N. Kampf, D. A. Jaitin, E. David, I. Amit, H. Nagase, I. Sagi, Proc. Natl. Acad. Sci. U. S. A. 2016, 113, 10884.

[26] J. Hazur, N. Endrizzi, D. W. Schubert, A. R. Boccaccini, B. Fabry, Biomater Sci 2021, 10, 270.

[27] O. Chaudhuri, L. Gu, M. Darnell, D. Klumpers, S. A. Bencherif, J. C. Weaver, N. Huebsch, D. J. Mooney, Nat. Commun. 2015, 6, 6364.

[28] Acta Biomater. 2019, 83, 221.

[29] S. A. Maskarinec, C. Franck, D. A. Tirrell, G. Ravichandran, Proc. Natl. Acad. Sci. U. S. A. 2009, 106, 22108.

[30] B. A. Nerger, M. J. Siedlik, C. M. Nelson, Cell. Mol. Life Sci. 2017, 74, 1819.

[31] J. Sapudom, S. Rubner, S. Martin, T. Kurth, S. Riedel, C. T. Mierke, T. Pompe, Biomaterials 2015, 52, 367.

[32] Y. M. Efremov, T. Okajima, A. Raman, Soft Matter 2020, 16, 64.

[33] Y. M. Efremov, W.-H. Wang, S. D. Hardy, R. L. Geahlen, A. Raman, Sci. Rep. 2017, 7, 1541.

[34] F. Charbonier, D. Indana, O. Chaudhuri, Curr Protoc 2021, 1, e124.

[35] M. Darnell, S. Young, L. Gu, N. Shah, E. Lippens, J. Weaver, G. Duda, D. Mooney, Adv. Healthc. Mater. 2017, 6, DOI 10.1002/adhm.201601185.

[36] C. Yang, F. W. DelRio, H. Ma, A. R. Killaars, L. P. Basta, K. A. Kyburz, K. S. Anseth, Proc. Natl. Acad. Sci. U. S. A. 2016, 113, E4439.

[37] K. Mandal, Z. Gong, A. Rylander, V. B. Shenoy, P. A. Janmey, Biomater Sci 2020, 8, 1316.

[38] J. Comelles, V. Fernández-Majada, N. Berlanga-Navarro, V. Acevedo, K. Paszkowska, E. Martínez, Biofabrication 2020, 12, 025023.

[39] A. Bauer, L. Gu, B. Kwee, W. A. Li, M. Dellacherie, A. D. Celiz, D. J. Mooney, Acta Biomater. 2017, 62, 82.

[40] M. Hörning, M. Nakahata, P. Linke, A. Yamamoto, M. Veschgini, S. Kaufmann, Y. Takashima, A. Harada, M. Tanaka, Sci. Rep. 2017, 7, 7660.

[41] J. L. Sacco, Z. T. Vaneman, E. W. Gomez, J. Cell. Physiol. 2024, 239, e31165.

[42] J. Y. Lee, A. A. Dominguez, S. Nam, R. S. Stowers, L. S. Qi, O. Chaudhuri, Sci. Rep. 2019, 9, 17188.

[43] J. Alcaraz, H. Mori, C. M. Ghajar, D. Brownfield, R. Galgoczy, M. J. Bissell, Integr. Biol. 2011, 3, 1153.

[44] E. E. Charrier, K. Pogoda, R. G. Wells, P. A. Janmey, Nat. Commun. 2018, 9, 449.

[45] A. Elosegui-Artola, A. Gupta, A. J. Najibi, B. R. Seo, R. Garry, C. M. Tringides, I. de Lázaro, M. Darnell, W. Gu, Q. Zhou, D. A. Weitz, L. Mahadevan, D. J. Mooney, Nat. Mater. 2023, 22, 117.

[46] W. Fan, K. Adebowale, L. Váncza, Y. Li, M. F. Rabbi, K. Kunimoto, D. Chen, G. Mozes, D. K.-C. Chiu, Y. Li, J. Tao, Y. Wei, N. Adeniji, R. L. Brunsing, R. Dhanasekaran, A. Singhi, D. Geller, S. H. Lo, L. Hodgson, E. G. Engleman, G. W. Charville, V. Charu, S. P. Monga, T. Kim, R. G. Wells, O. Chaudhuri, N. J. Török, Nature 2024, 626, 635.

[47] M. S. Hall, F. Alisafaei, E. Ban, X. Feng, C.-Y. Hui, V. B. Shenoy, M. Wu, Proc. Natl. Acad. Sci. U. S. A. 2016, 113, 14043.

[48] S. Huth, S. Sindt, C. Selhuber-Unkel, PLoS One 2019, 14, e0220281.

[49] K. M. Schultz, K. S. Anseth, Soft Matter 2013, 9, 1570.

[50] C. B. Raub, J. Unruh, V. Suresh, T. Krasieva, T. Lindmo, E. Gratton, B. J. Tromberg, S. C. George, Biophys. J. 2008, 94, 2361.

[51] R. Murphy, A. Turcott, L. Banuelos, E. Dowey, B. Goodwin, K. O. Cardinal, Tissue Eng. Part C Methods 2020, 26, 628.

[52] C. E. Tolbert, L. Palmquist, H. L. Dixon, M. C. Srougi, J. Vis. Exp. 2019, DOI 10.3791/59989.

[53] U. Horzum, B. Ozdil, D. Pesen-Okvur, MethodsX 2014, 1, 56.

[54] J. Schindelin, I. Arganda-Carreras, E. Frise, V. Kaynig, M. Longair, T. Pietzsch, S. Preibisch, C. Rueden, S. Saalfeld, B. Schmid, J.-Y. Tinevez, D. J. White, V. Hartenstein, K. Eliceiri, P. Tomancak, A. Cardona, Nat. Methods 2012, 9, 676.

[55] J.-Y. Tinevez, N. Perry, J. Schindelin, G. M. Hoopes, G. D. Reynolds, E. Laplantine, S. Y. Bednarek, S. L. Shorte, K. W. Eliceiri, Methods 2017, 115, 80.

